# Contiguous Erosion of the Inactive X in Human Pluripotency Concludes With Global DNA Hypomethylation

**DOI:** 10.1101/2020.05.20.100891

**Authors:** Prakhar Bansal, Stefan F. Pinter

## Abstract

Female human pluripotent stem cells (hPSCs) are prone to undergoing X chromosome erosion (XCE), a progressive loss of key epigenetic features on the inactive X that initiates with repression of *XIST*, the long non-coding RNA required for X inactivation. As a result, previously silenced genes on the eroding X (Xe) reactivate, some of which are thought to provide selective advantages. To-date, the sporadic and progressive nature of XCE has largely obscured its scale, dynamics, and key transition events.

To address this knowledge gap, we performed an integrated analysis of DNA methylation (DNAme), chromatin accessibility, and gene expression across hundreds of hPSC samples. Differential methylation across the Xe enables ordering female hPSCs across a trajectory of XCE from initiation to terminal stages. Our results identify a crucial *cis*-regulatory element for *XIST* expression, trace contiguously growing domains of reactivation to a few euchromatic origins on the Xi, and indicate that the late-stage Xe impairs DNAme genome-wide. Surprisingly, from this altered epigenetic landscape emerge select features of naïve pluripotency, suggesting its link to X chromosome dosage may be partially conserved in human embryonic development.

## INTRODUCTION

Human pluripotent stem cells (hPSCs) of embryonic (ESC) or induced (iPSC) origin resemble the inner cell mass (ICM) of the early post-implantation embryo. Capable of differentiating into all three germ layers, hPSCs have become a corner-stone for modeling human disease *in vitro*^1^, and have tremendous potential for future allogenic or autologous cell replacement therapies^2^. However, in addition to previously noted challenges to their genetic^3–6^ and epigenetic^7–9^ stability, female-derived hPSCs in particular may be expected to lag behind male hPSCs in some applications due to progressive reactivation of their inactive X (Xi), a process termed X chromosome erosion (XCE)^10–12^.

In contrast to female hPSCs that reflect a late epiblast (or “primed”) cell state after completed X chromosome inactivation (XCI), female ESCs and iPSCs of the mouse approximate the ICM of the (“naïve”) pre-implantation embryo, and harbor two active X (Xa) chromosomes^13^. To transition from naïve to primed pluripotency and differentiate, mouse ESCs must silence all supernumerary X chromosomes, to maintain only a single Xa per diploid genome^14–16^. This obligate coupling of pluripotency state to X dosage has propelled most of our current understanding of random X chromosome inactivation (XCI)^17,18^. Yet, recent studies of peri-implantation embryos across a range of mammals have shed light on important species-specific differences in XCI establishment^19–23^.

XCE in female hPSCs is an *in-vitro*-and species-specific defect in XCI maintenance that is best understood as two mechanistically distinct observations. First, post-XCI mouse ESC-derived epiblast cells faithfully maintain *Xist* expression and gene silencing of the Xi^23^, whereas female primed hPSCs show a strong tendency to lose *XIST* expression^24,25^. Second, even upon deletion of *Xist*, the vast majority of genes on the mouse Xi remain stably silenced on account of long-term and selfpropagating DNA methylation (DNAme) of promoters and CpG islands^26–28^, with rare and context-specific exceptions^29–31^. Moreover, even a brief period (< 3 days) of mouse *Xist* expression is sufficient to irreversibly silence genes long-term^32^. In contrast, female hPSCs that have lost *XIST* expression have been demonstrated to undergo frequent, progressive and irreversible X reactivation^10,33–37^.

Yet, despite the relevance of hPSCs to X-linked disease modeling and future cell replacement therapies, our mechanistic understanding of XCE has remained limited, with most studies confined to a small number of lines. As a result, descriptions of XCE have ranged from minor^24^, major^38^ to whole Xi reactivation^10^, indicating that the progressive nature of this phenomenon may require analysis across a larger number of female hPSC lines. Additional observations from gene expression analyses contrasting ESC vs. iPSC lines^39^, or preferential reactivation of X-linked oncogenes^36^, have not been mirrored in larger DNAme analyses^10,38^.

To address this knowledge gap, we performed an integrated analysis of available DNAme, chromatin accessibility and gene expression data across 399 hPSC lines. Differential DNAme across the eroding X (Xe) orders female hPSCs across a trajectory of XCE from initiation to terminal stages. We find that XCE spreads in contiguous fashion from a few euchromatic regions that escape XCI, and observe that terminal XCE results in female-specific DNA hypomethylation that triggers some markers of naïve pluripotency. Our analysis is relevant to understanding and staging the epigenetic fidelity of female hPSC disease models, and reveal the impact of X chromosome dosage on embryonic stagespecific pluripotency.

## RESULTS

### Female hPSC samples trace a common path through XCE

To assemble a comprehensive view of X chromosome methylation dynamics in human pluripotency, we collated Illumina BeadChip (450K, MethylEpic) data from all suitable hPSC studies in GEO^38,40–46^. After removal of hPSC lines with incorrectly listed sex chromosomes (see methods), the resulting superset spanned 471 hPSC lines (Fig. S1A). Because X-linked DNAme (β-values) in female hPSCs are averaged between Xi and Xa, probes in genes subject to XCI center around a β-value of 0.5 (Fig. 1A). Male hPSC lines cluster together in PCA (Fig. 1B), where PC1 reflects the large range of average X chromosome DNAme of female hPSCs (Fig. S1B-E). There was little to no influence from the originating study, hESC/iPSC category or hPSC identifier, on either PC, except for an over-represented cluster of the predominant H9 hESC model that skewed overall analysis and was removed (see methods). This balanced superset of 178 male and 221 remaining female hPSC lines revealed excess female-specific variance in X-linked probes, with no overall sex differential across autosomal probes (Fig. 1C,D). This confirms that XCE cannot result from genome-wide DNAme differences between the sexes^38^, and enables identifying XCE-relevant probes on the basis of their significant excess variance across female relative to male hPSCs (Fig. S1F,G). Indeed, significantly-varying probes (p ≤ 0.01, Brown-Forsythe test) on the X have a bimodal distribution that reflects genes subject to XCI (centered around 0.5), and likely escapee genes that are hypomethylated on both Xa and Xi, with a wide range of probes between these two peaks (Fig. S1H). Average X DNAme in female hPSCs correlates almost perfectly with the sex-variant probes, exceeding its correlations with non-variant X-linked probes or genome-wide DNAme levels (Fig. S1I,J).

**Figure 1:**
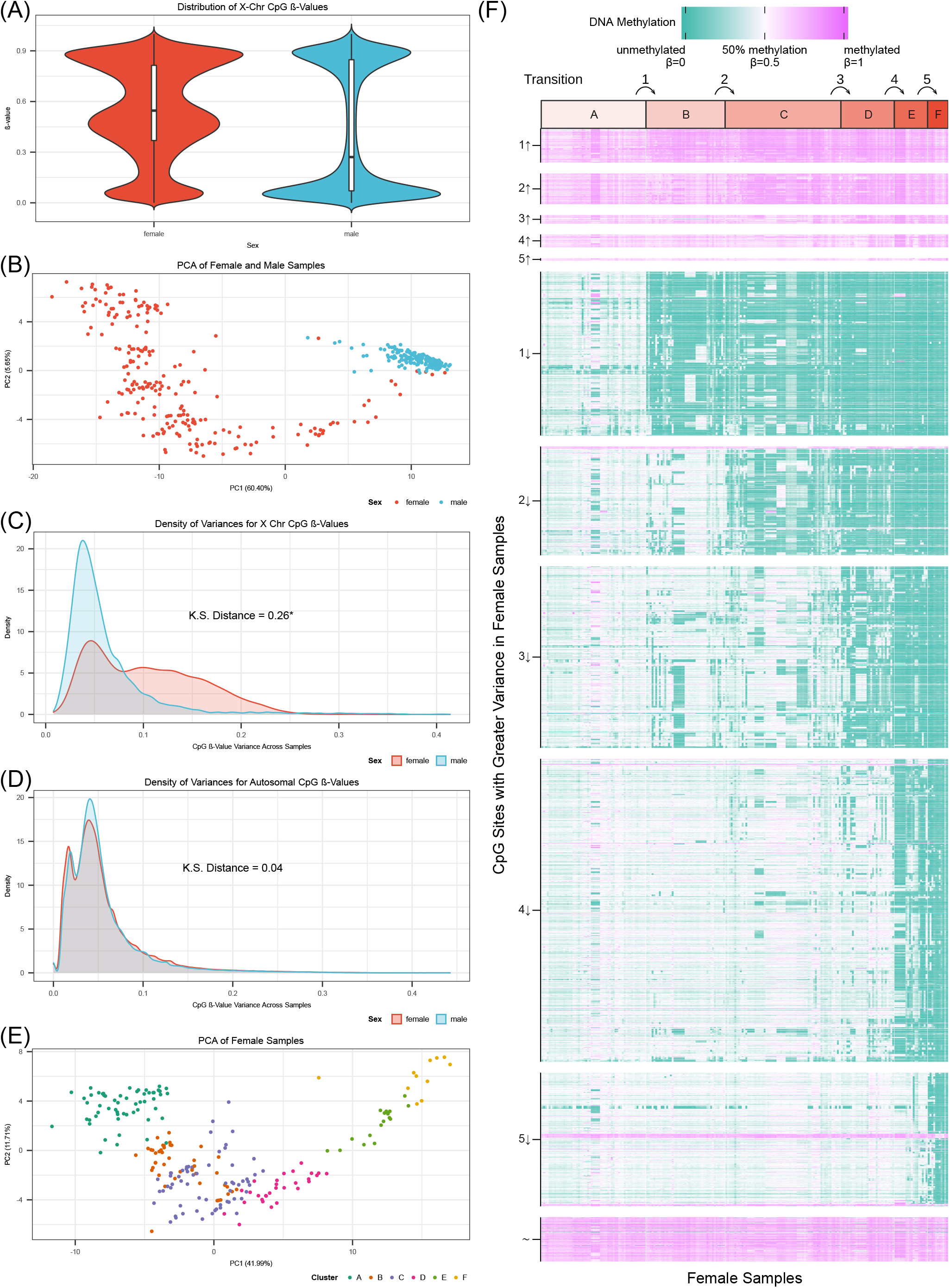
XCE reflected in high X-specific variance in female hPSCs. (A) ß-values distribution of probes on the X in female (red) and male (blue) samples. (B) Principal component analysis (PCA) of female (red) and male (blue) samples using only X-linked probes. (C) and (D) Density plots of probe β variance on X and autosomes, respectively, with red lines for female samples, and blue for male samples (Kolmogorov-Smirnov, KS distance indicated, with KS p-values of 5.6e-15 and 0.78 for C and D, respectively). (E) PCA of female samples colored by their k-means assigned cluster (see methods). (F) DNAme heatmap of high variance X probes in female hPSCs. Samples within each k-means cluster (A-F) are ordered by mean X DNAme. Probes (rows) are grouped by transition (lowest q-value ≤ .05), and β-value change (↑ up, down ↓, or fluctuating irrespective of transition ~). Cluster transitions are numbered (1-5).

To stage XCE progression, we used probes with high sex-variance to perform K-means clustering of female samples, yielding six clusters (Fig. 1E, Fig. S1K) ordered by their average X DNAme. We then identified all differentially methylated probes (DMPs) between neighboring clusters (Fig. 1F). Despite inclusion of both iPSCs and ESCs from eight independent studies, DNAme is remarkably similar for any given probe within each cluster, demonstrating that most reactivation events are not sporadic. Moreover, of nine hPSC lines contributing to more than a single cluster, eight revealed passage number increases consistent with this cluster order, suggesting that they reflect a common XCE timeline. Overall, we see a clear, systematic pattern of DNAme changes (Fig. 1F) that illustrates stepwise XCE progression. Probes losing DNAme far outnumber probes gaining DNAme, with promoter-associated probes pre-dominating amongst de-methylated DMPs (Fig. S2A). Of 450 X-linked genes that are subject to XCI (promoter-associated β between 0.3-0.7 in cluster A), only seven (1.6%) maintain their DNAme in cluster F, suggesting that gene reactivation across the Xe is virtually complete.

### *XIST* and the *FIRRE* tandem repeat undergo hypermethylation at the onset of XCE

With this progressive staging of XCE in hand, we parsed step-wise DNAme changes occurring at each transition between clusters A-F. Analysis of transition 1 (cluster A to B) reveals that DMPs reside in specific regions (Fig. 2A, blue line), in a pattern that matches neither overall probe density (grey, 9191 probes), nor the 1412 probes that fail the DMP threshold (q>0.05, black line, Fig. S2B). DMP densities of decreasing DNAme generally match those with increasing DNAme, indicative of regional Xi gene reactivation, with lower and higher DNAme of promoter-associated versus transcribed intragenic CpGs, respectively.

**Figure 2:**
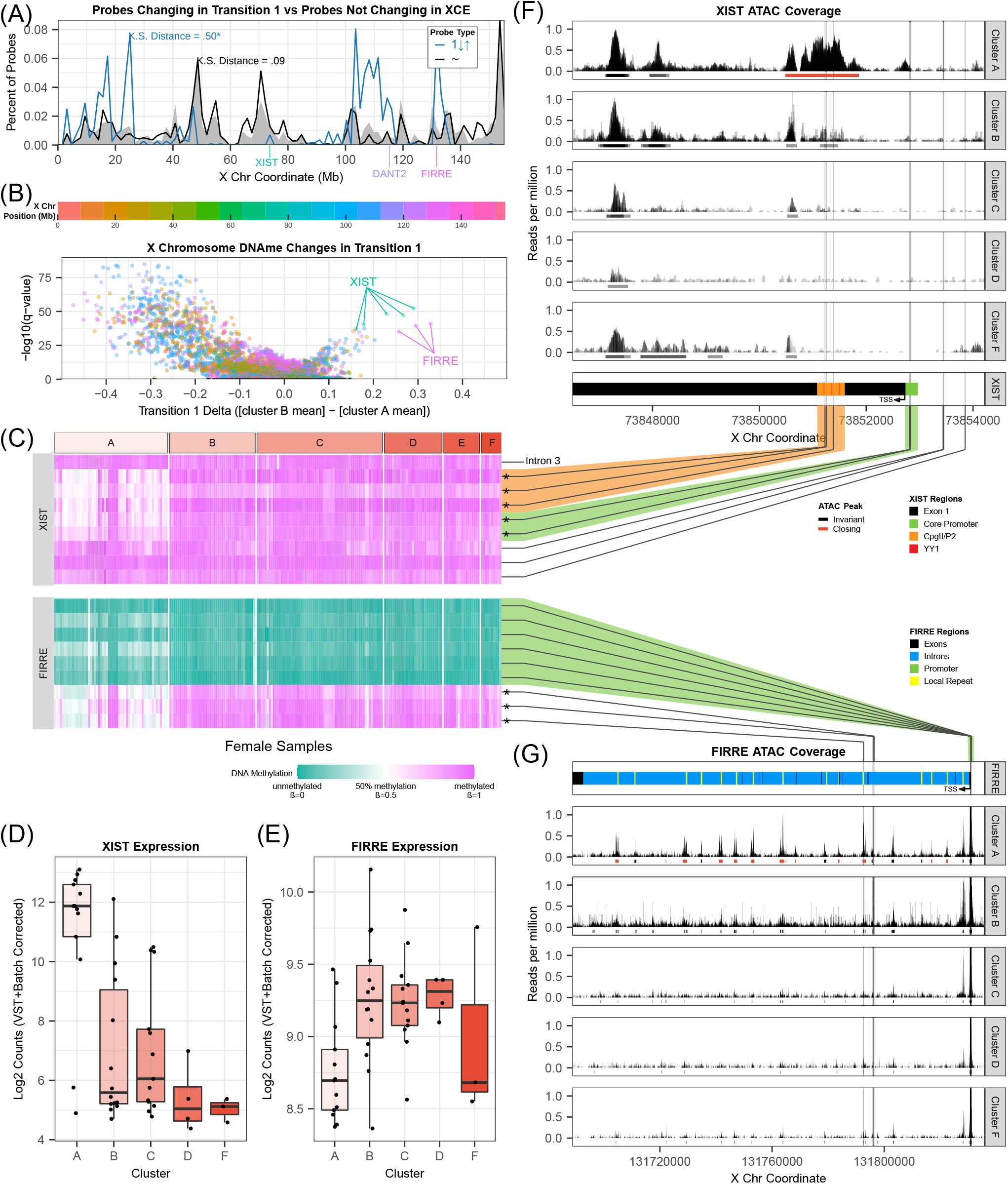
Differential DNAme analysis on Xe distinguishes early from late XCE. (A) Density plot of probes across X coordinates reveal transition 1 probes (1↓↑, blue) to concentrate irrespective of overall probe density (grey shaded area), in contrast to fluctuating probes (~, black). K.S. distances quantify similarity to frequency distribution of the null probe density distribution (* K.S. test p-value=5.09e-11 for blue transition 1 probes). (B) Volcano plot of transition 1 differentially methylated probes (DMPs), colored by X coordinate (legend above the plot). Top increasing DMPs annotated by gene name. (C) DNAme of *XIST* and *FIRRE* probes across clusters A-F. Lines to (F) and (G) indicate position of probes in relation to the genes and their annotated regions (* indicates DMPs from B). (D) and (E) *XIST* and *FIRRE* expression (VST+batch-corrected/log2-transformed), respectively, for all samples with available RNA-seq data (all but cluster E). (F) and (G) ATAC-seq coverage for *XIST* and *FIRRE*, respectively. Horizontal lines below coverage indicate called peaks (gray, black), including differential, closing peaks (red). Bottom panels of (F) and (G) relate DMPs and ATAC-peaks to *cis* regulatory sites for *XIST* and *FIRRE*, respectively.

For transition 1, eight DMPs stand out in degree and significance of increasing DNAme (Fig. 2B,C), with five DMPs in *XIST* and the three in the tandem repeat of *FIRRE*. Loss of XIST expression has been suggested as an early or causative event for XCE^10,24,25,37,47^, but its mechanistic basis has remained unclear. Out of the 9 probes spanning *XIST*, five change in transition 1, two of which are located in its core promoter^48^. The other three DMPs are located in a likely enhancer in exon 1 (~ 1.2 kb from the *XIST* promoter), two of which directly overlap binding sites for the YY1 transcription factor (Fig. 2C,F). The YY1 cognate motif is conserved across a range of mammalian *XIST* homologues, and YY1 is necessary for *Xist* expression in mouse ESCs^49^. YY1 binding is sensitive to DNAme^50^, and in differentiated cells, only the silent *XIST* allele on the Xa features methylated YY1 cognate motifs^51^. Indeed, as these same YY1 sites gain DNAme in transition 1, the major overlapping ATAC peak is lost and *XIST* expression drops by over two orders of magnitude (Fig. 2F,D).

Three other top-ranking DMPs increasing in DNAme map to the *FIRRE* locus, which is also the only X-linked gene outside of *XIST* to lose ATAC peaks in transition 1 (Fig. 2C,G). This evolutionarily conserved tandem repeat forms long-range CTCF-mediated chromatin loops with two other tandem repeats *DXZ4* and *ICCE* on the Xi, and transcribes a long non-coding RNA (ncRNA) primarily from the Xa^52,53^. The three *FIRRE* DMPs gaining DNAme reside inside the tandem repeat, characterized by closing (“local repeat”) ATAC peaks, while probes near its major promoter decrease in DNAme (Fig. 2C,G). This is consistent with gene reactivation, although the modest increase in *FIRRE* expression was not statistically significant (Fig. 2E), and may reflect only partial escape from XCI, or an escapee-like chromatin state as seen for mouse *Firre*^54,55^. The respective gain and loss of DNAme and chromatin accessibility of tandem repeat elements inside *FIRRE* mirror changes at *XIST* in transition 1, and suggest that this conserved ncRNA gene is among the very first to reflect XCE.

### Contiguous progression of XCE originates near escapee genes

While the *XIST*-neighborhood appeared to remain refractory to early loss of DNAme (Fig. 2A), genes near *FIRRE (*130 Mb, magenta), *XACT* and *DXZ4* (110-115 Mb, blue/lavender) on the long arm of the X, and on the escapee-rich short arm, experience the earliest and most significant loss of DNAme in XCE. Tracking transition-specific significant DMPs (Fig. 3A, colors) and cumulative DNAme changes (grey), we find that XCE appears to emanate from these specific chromosomal regions, and then spreads in contiguous fashion across the rest of the X chromosome during subsequent transitions (Fig. 3A). This wave of DNAme loss is mirrored by progressive (~2-fold) gains in chromatin accessibility in the same regions (Fig. 3B & inset). Curiously, DMPs and opening ATAC peaks specific to early transitions reside in H3K27me3-rich regions of the Xi, which are likely most sensitive to the loss of *XIST* RNA, while those of later transitions tend to fall in H3K9me3-rich regions. Such association with specific chromatin compartments is best reflected in the progressive shift of DMP and differential ATAC peak density correlating with H3K27me3 levels to H3K9me3 in late XCE (Fig. 3C,F). In sum, these DNAme and chromatin accessibility patterns strongly suggest that XCE is dictated by linear spreading of reactivation along the chromosome.

**Figure 3:**
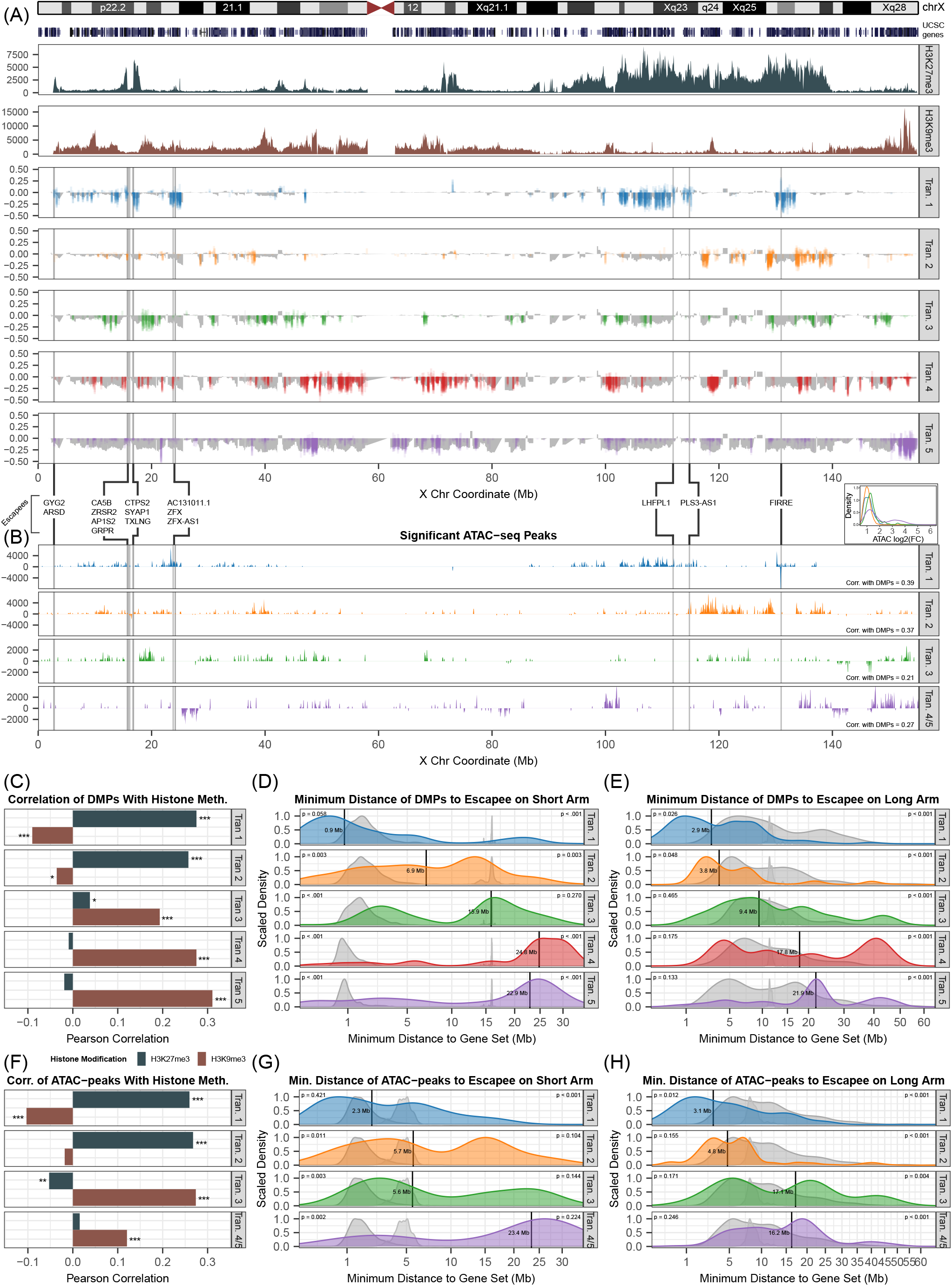
Contiguous spread of XCE from euchromatic origins on the Xi. (A) Map of cumulative (grey) and transition-specific DMP (colors) changes in β-values across the X plotted underneath Xi-specific H3K27me3 (dark teal) and H3K9me3 (brown) H9 ChIP-seq from^37^. Grey vertical lines indicate escapee locations, determined from DNAme in cluster A, as in^57^. (B) Map of coverage of differential ATAC-peaks across the X. Negative coverage indicates a negative fold change. Transitions 4 and 5 are combined (purple), due to lack ATAC-seq data for cluster E. Histograms of log2FC of differential peaks for each transition are inset on top right (inset). (C) Pearson correlations of transition-specific DMP changes (colored in A) to H3K27me3 (dark teal) and H3K9me3 levels (brown). (D) and (E) DMP distance distributions relative to escapee locations for each transition on short arm and long arm, respectively, with black vertical lines denoting median distance. Null distance distributions using permuted escapees (broad grey) or permuted DMPs (narrow grey) plotted underneath actual distributions. Calculated p-values (two-tailed rank test against the permutated distributions) indicated on the top-left (permuted escapees) and top-right (permuted DMPs) of each plot. (F) Pearson correlations of transition-specific ATAC-peaks (B) to H3K27me3 (dark teal) and H3K9me3 levels (brown). (G) and (H) ATAC-peak distance distributions relative to escapee locations for each transition on short arm and long arm, respectively, with black vertical lines denoting median distance.

In tracing such contiguous XCE to its origins, we noticed that early DMPs and opening ATAC peaks map near genes known to escape XCI, both on the escapee-rich short arm, and on the long arm from two regions in particular (Fig. 3A,B): 1.) the ~5-Mb focal domain centered on *FIRRE*, and 2.) the ~15-Mb wide domain to the left of the *DXZ4* macrosatellite (near *PLS3-AS1*), previously shown to require *DXZ4* for H3K27me3 maintenance^56^. To confirm and characterize this pattern quantitatively, we first identified *bona fide* escapee genes in hPSC lines of cluster A using the approach by Cotton, et al.^57^. Altogether we identified 15 genes that, based on DNAme, are likely to escape XCI in non-eroding, *XIST*-positive hPSCs (Fig, S2E-G). In stark contrast to genes subject to XCI, these genes feature largely hypomethylated promoters and hypermethylated gene-bodies due to their bi-allelic expression (Fig. S2H). Additionally, pseudo-autosomal region 1 (PAR1) is not represented on Illumina beadchips, but known to escape XCI. We therefore tested whether the minimal linear distance from short-arm/PAR1 or long-arm escapees, segregated DMPs or differential ATAC peaks by their transition (Fig. S2I-N). Indeed, these cumulative distributions suggest that XCE traverses between ~3-9 Mb for most transitions, with a median escapee distance of ~15 Mb (DNAme) and ~10 Mb (ATAC) at the XCE mid-point in transition 3, and reaching H3K9me3 chromatin last in transitions 4 and 5 (Fig. 3A-C,F).

To determine the significance of this observation, we performed two permutation tests for each transition. In the first, we randomized the 15 escapee labels across all genes represented by probes, while in the second test we randomized the transition labels for each DMP/ATAC peak. The resulting distributions of these permutations are represented as broad (randomized escapees) and sharp (randomized DMPs/ATAC peaks) grey background distributions (Fig. 3D,E,G-H). Indeed, on both long and short arms, DMPs and opening ATAC peaks associated with transition 1 (blue distribution) are significantly closer to the 15 escapees than at least one of their randomized permutation sets. At subsequent transitions, the real median DMP/ATAC peak-escapee distance progressively approaches and eventually surpasses both randomized distance sets. In sum, the progressively increasing linear distance of DMPs and ATAC peaks to escapees serves as a strong predictor for which genes are most likely to undergo XCE early vs. late, which suggests that XCE is likely to spread in contiguous fashion.

### Terminal XCE is associated with global DNA hypomethylation

While the first three transitions primarily impacted the X, genome-wide DNAme and expression analysis reveals an emergent and dominant *trans*-effect as XCE nears completion: relative to clusters A-D, hPSC samples of cluster F show significant autosomal hypomethylation compared to other clusters and male samples (Fig. 4A), This global reduction of DNAme is paralleled by biased upregulation of autosomal genes (Fig. 4 B,C). Such global effects are particularly striking when compared to relatively X-specific changes occurring at transitions 1-3, with some early loss of autosomal DNAme starting in transition 4, and rapidly amplifying in transition 5 (Fig. S3), when a plethora of DMPs across the genome drop to reduce cumulative DNAme.

**Figure 4:**
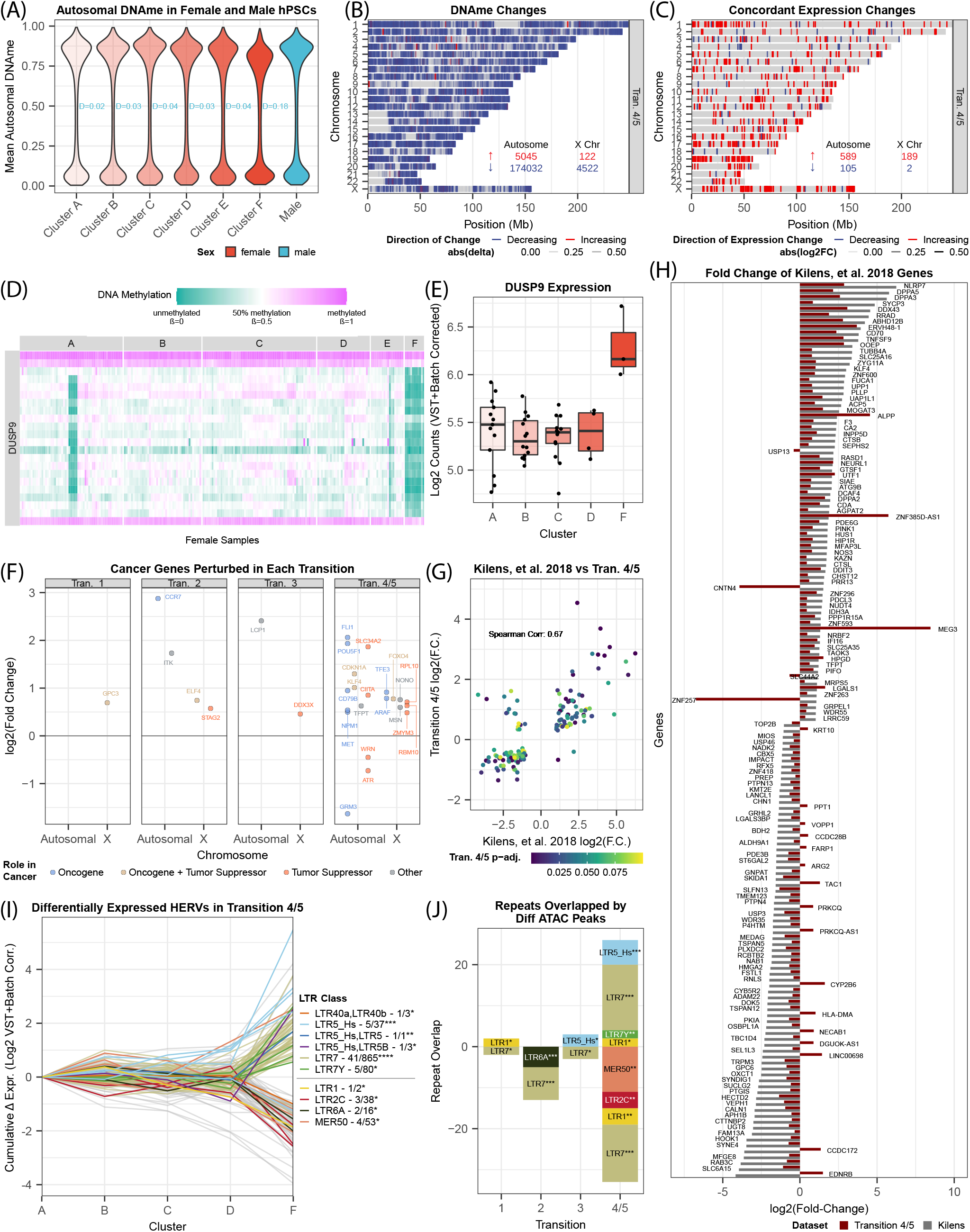
Global hypomethylation and emergent naïve pluripotency markers in terminal XCE. (A) Autosomal methylation distributions across female hPSC clusters (A-F, red shades) and their K.S. distances relative to male hPSC distribution (teal). (B) Chromosome-resolved DMPs of the final XCE transitions (4 or 5). Total number of probes changing for autosomes and X listed with colors corresponding for increasing (red) or decreasing (blue) DNAme. The transparency is on a continuous scale based on the β-differential. (C) As (B) but plotting differentially expressed genes (DESeq2) with concordant DNAme changes (B) and/or ATAC changes. The transparency is on a continuous scale reflecting the log2(fold change). (D) DNAme heatmap of *DUSP9* probes across clusters A-F. (E) *DUSP9* (VST+batch-corrected/log2-transformed) expression across clusters A-F. (F) Transition -and autosome/X -resolved differentially expressed genes annotated in the COSMIC cancer gene census^79^. (G) Differential expression scatter plot of transition 4/5-specific genes (Y-axis) over log2-fold changes (X-axis) of these genes between naïve and primed hPSCs from^64^. (H) Comparison of expression changes in transition 4/5-specific differential genes (dark red) relative to corresponding changes (grey) in naïve vs. primed hPSCs from^64^. (I) Cumulative changes (relative to cluster A) in all differentially expressed HERVs identified by Telescope^67^ (VST+batch-corrected/log2-transformed). Only significantly overrepresented HERVs (hypergeometric p ≤ 0.1) are labeled by colors (other in grey). Color legend lists total counts of differentially expressed HERV classes (with hypergeometric significance indicated as * < .1, ** < .05, ***<.01, ****<.001). (J) Tally of differentially expressed HERV classes (I) overlapped by significantly differential ATAC peaks (fisher right-side p-value indicated as *<.01, ** <.001, ***<.0001). Negative overlaps indicate overlaps by closing ATAC peaks, and positive by opening peaks.

In principle, two competing hypotheses for this effect can be considered: 1.) cluster F may comprise of aberrant hPSC lines that suffer some generalized loss of facultative heterochromatin, or 2.) terminal XCE specifically reduces global DNAme, likely via overexpression of X-linked genes that impact DNAme *in trans*. Arguing against the former, overall autosomal chromatin accessibility drops in transition 4 with far fewer opening than closing peaks (Fig. S3), which primarily reside in sparsely accessible non-coding regions and near cell-type specific clustered paralogous gene families (e.g. olfactory receptors, keratins, cadherins). In contrast, and in support of the latter hypothesis, genomewide hypomethylation of XaXa mouse ESCs relative to 39,X and 40,XY cells has previously been linked to double-copy dosage of the X-linked *Dusp9* gene^58^. Consistent with this observation, cluster F hPSC samples uniformly show loss of DNAme at the *DUSP9* promoter, paralleled by *DUSP9* overexpression in this last transition (Fig. 4D, E). Indeed, autosomal DNAme correlates strongly with *DUSP9* methylation in male and female hPSCs, but the slope of this relationship steepens in the context of two active *DUSP9* copies in cluster F hPSCs (Fig. S4A). Expression of mouse *Dusp9* from two active copies in XaXa ESCs has been shown to impact global DNAme by inhibiting the MAPK pathway, resulting in a post-translational decrease in DNMT3A/B and UHRF1 proteins that was not evident in *Dnmt3a/b* and *Uhrf1* transcript levels^58^. Likewise, neither human orthologue was differentially expressed in hPSC samples of cluster F (not shown). Two additional factors signaling via the MAPK pathway (*ELK1*^59^ and *ARAF*) reside in another H3K9me3-rich region of the hPSC Xi, and increase ~2-fold upon loss of promoter-associated DNAme (Fig. S4C,D). We also tested whether oncogenes are preferentially activated^10,47^, but find only a handful of *bona fide* oncogenes specific to the last transition on X (*ARAF, TFE3*) and autosomes (*MET, NPM1, CD79B, POUF5F1 (OCT4), FLI1*) alike (Fig. 4F). Consistently, there was only a single cancer-associated term enriched among X-linked genes differentially expressed in transition 5, undermining the notion that oncogene activation provides a dominant selective benefit for hPSCs undergoing early XCE.

While MAPK inhibition is essential for *in vitro* induction of naïve pluripotency^60,61^ it also abrogates global DNAme and erases imprinting^62^. Motivated by the observation that reduced inhibition of MAPK signaling is sufficient in naïve pluripotency^63^, we tested whether the modest attenuation of DNAme levels in cluster F hPSCs is reflected in expression differences of their pluripotency genes. To this end, we compared gene expression profiles of naïve and primed hPSCs derived in parallel^64^ to genes differentially expressed in cluster F. The overlap (189/1686) and correlation (rho=0.67, p=2.08e-21) across differentially expressed genes greatly exceeded our expectations (Fig. 4G,H), with naïve pluripotency factors *DPPA2/3/5* and *KLF4* upregulated alongside Kelch homology (KH) or KH domain-like genes *NRLP7* and *OOEP* (also *DPPA5*). Moreover, among upregulated genes we find many specifically expressed in three embryonic pre-implantation stages (zygote, morula and blastocyst), including *DDIT3, NANOG, POU5F1 (OCT4), TLE6* and *ZSCAN4*, among many others. Upregulation of these genes is significantly reflected in reduced DNAme (Fig. S4B, sign test p < 2.2e-16). Because the overall degree of these changes lagged behind the dynamic range observed in the bona fide naïve hPSCs, we also examined whether there were changes in human endogenous retroviral (HERV) elements, which are expressed in stage-specific fashion during mammalian pre-implantation development and feature pluripotency factor binding sites in their long terminal repeats (LTRs)^65,66^. Indeed, retrotranscriptome analysis using Telescope^67^ reveals significantly-enriched upregulation of HERV-H (LTR7Y p=2.7e-4) and HERV-K (LTR5_Hs p=0.003) elements, which are respectively associated with blastocyst and morula-stage embryos^68^ and likewise upregulated in naïve hPSCs in vitro^60,63,66^. While these HERV classes were also significantly enriched for opening ATAC peaks in cluster F (with 16 LTR7, two LTR7Y and six LTR5_Hs elements), no naïve pluripotency-specific upregulation or chromatin opening of hominid SVA elements^60,69^ was observed in these hPSCs (Fig. 4J). In sum, these changes suggest that despite their more limited hypomethylation relative to *bona fide* naïve hPSCs (Fig. 4H), cluster F hPSCs appear to regain some specific hallmarks of naïve pluripotency.

## DISCUSSION

Our integrated DNAme, ATAC and expression analysis addresses the three major challenges that have obscured our understanding of XCE in female hPSC to-dates: 1.) an incomplete view of human *XIST* regulation, 2.) the progressive nature of XCE coupled with a lack of longitudinal studies, and 3.) a consequently large range of reported outcomes, from piecemeal to whole Xi reactivation^10,33–38,70^. In ordering over 200 female hPSC samples from early to late XCE, this analysis leads us to four novel insights regarding this elusive epigenetic phenomenon.

First, we pinpoint a dynamic increase of DNAme and drop in chromatin accessibility in a *cis-*regulatory CpG island just 1.2 kb downstream of *XIST’s* major promoter, that coincides with loss of *XIST* expression and the very earliest Xi reactivation events (Fig. 2). The same “P2” element is one of only three female-specific DNase hypersensitive sites on the human X, and is fully methylated on the single Xa in male cells^51,71^. The corresponding DMPs overlap conserved cognate motifs of transcription factor YY1 that binds in DNAme-sensitive manner to boost both human and mouse *XIST/Xist* expression^49^. Two equally parsimonious causes for loss of *XIST* expression could be that continued selection for high expression of pluripotency factors, including REX1/ZFP42 that directly competes for binding the YY1 motif^49^, or aberrant MYC-mediated DNAme in hPSCs^7^, result in YY1 eviction from the P2 element. Indeed, REX1/ZFP42 is required in human pluripotency^72^, and joins another pluripotency factor (*KLF4*) known to repress Xist in the mouse^73^, in progressively increasing expression across XCE (Fig. S2C,D). Of the two remaining female-specific YY1 clusters on the human X^51,71^, *FIRRE’s* tandem repeat also reflects this first XCE transition with some of the most significantly increasing DMPs and the only closing ATAC peaks on the X outside of *XIST* (Fig. 2,3). *FIRRE* has been shown to form conserved Xi-specific long-range interactions with the third female-specific YY1 cluster, the macrosatellite *DXZ4* which functions as a topological boundary on the Xi (reviewed in^53^). Both of these non-coding tandem repeat loci maintain focal euchromatin on the otherwise escapee-poor long arm, and reside at the center (*FIRRE*) or periphery (*DXZ4*) of the earliest reactivation clusters (Fig. 3). Interestingly, the DXZ4-adjacent 15-Mb region was shown to lose H3K27me3 when *DXZ4* is deleted in differentiated RPE-1 cells^56^. This region also spans *XACT*, another ncRNA gene associated with XCE initiation in H9 hESCs^37^.

Second, in addition to H3K27me3/H3K9me3 association, we find that the linear distance from escapee genes is a major predictive factor for whether a given gene loses DNAme and becomes more accessible during early or late XCE (Fig. 3). To this end, we identified 15 escapee candidates in female hPSCs on the basis of their DNAme states, as in Cotton, et al.^57^. On the long arm, novel escapee candidate *LHFPL1* (~ 1 Mb from *XACT*), joined *PLS3-AS1* (~ 0.25 Mb from *DXZ4*), and *FIRRE*, while on the short arm, outside PAR1, twelve genes qualified as escapees based on their DNAme. On both long and short arm, DMPs associated with later transitions increased in distance from these escapee candidates, and this contiguous spread was also evident in progressive gains of chromatin accessibility (Fig. 3A,B). This pattern resembles the collinear activation of autosomal HOX clusters, which are repressed by H3K27me3 chromatin and progressively activated towards the embryonic posterior^74^. While we cannot demonstrate causality, one plausible mechanistic basis for this pattern is that active chromatin spreads from escapee genes to reactivate silenced genes in contiguous fashion. In support of this notion, deletion of an escapee boundary does not abrogate its expression on the mouse Xi, but instead drives reactivation of neighboring genes^75^. In another X reactivation context during mouse iPSC reprogramming, genes closest to escapees also reactivate before distal genes^76^.

Third, terminal XCE is strikingly associated with female-specific global DNA hypomethylation (Fig. 4A,B,C), as previously demonstrated in XaXa mouse ESCs^58^. As in the mouse system, we find that doubling of *DUSP9* dosage, which is specific to the final XCE transition, correlates with autosomal hypomethylation (Fig. S4A). We find little evidence for preferential upregulation of oncogenes before terminal XCE, when two X-linked and five autosomal oncogenes are activated (Fig. 4F). We therefore conclude that selection for female hPSCs undergoing XCE is unlikely to drive early XCE. Surprisingly, in terminal XCE, when reactivation of *DUSP9* impairs the MAPK pathway and DNAme genome-wide is reduced, we find a highly significant shift towards a naïve pluripotent gene expression profile, as primed markers are repressed and naïve markers are induced (Fig. 4G,H). While the magnitude of naïve marker induction lags behind that observed for *bona fide* naïve hPSCs, reactivated and reopening HERVs (Fig. 4I, J) match the *in vivo* equivalent preimplantation-stage specific HERV classes, namely HERV-H (blastocyst) and HERV-K (morula)^68^. This HERV profile is also activated either transiently during iPSC reprogramming^77^, or stably upon induction naïve pluripotency^60,63^. In mouse ESCs, double X chromosome dosage blocks exit from naïve pluripotency^16^, likely by stabilizing expression of specific pluripotency regulators^78^. As a result, pluripotency exit is coupled to XCI. We therefore interpret the basal induction of naïve-specific genes and HERVs in terminal XCE (Fig. 4) as hPSCs partially recapitulating this process in reverse, suggesting the conservation of this double X dosage impact on the human naïve pluripotency circuit. Intriguingly, enrichment of upregulated mitochondrial factors, and regulators oxidative phosphorylation in terminal XCE, may hint at a possible link to a third hallmark of naïve pluripotency, namely increased mitochondrial respiration^64^.

Looking forward, this analysis raises important questions for future work regarding possible roles of REX1/ZFP42 or aberrant DNAme in evicting YY1 from *XIST’s* P2 element, and whether *FIRRE* and *DXZ4* are sensitive to the same pluripotency circuit, or respond to the earliest Xi structural changes upon loss of *XIST*. Moreover, if euchromatic regions indeed seed X reactivation, is it possible that *FIRRE* and *DXZ4* have been conserved in evolution due to the paucity of escapee genes on the long arm of the human X? Although neither of their mouse orthologues is required for XCI *in vivo*, their potential roles in X reactivation remain to be addressed in detail. The high density of escapees and the active PAR1 may also predispose the short arm of the human Xi to XCE, which may explain why hPSCs depend on *XIST* for maintenance of XCI, in contrast to the mouse system. In view of potential application of hPSC-derived cells for disease modeling and autologous cell replacement therapies, it would also be important to ascertain whether early or intermediate XCE halts upon differentiation, or progresses thereafter. Finally, identifying which X-linked factors apart from *DUSP9* may be necessary and sufficient to trigger female-specific hypomethylation in hPSCs represents a crucial goal towards understanding why the female/Xa-dosage bias towards naïve pluripotency may have been conserved in evolution^16,78^.

## ACKNOWLEDGMENTS

We thank the lab for critical reading of the manuscript. This work was supported by NIH grant R35GM123926 to S.F.P.

## AUTHOR CONTRIBUTIONS

P.B. collated, processed and analyzed all data. P.B. and S.F.P. conceived of the study, interpreted the results and wrote the manuscript.

## DECLARATION OF INTERESTS

No competing interests

## METHODS

### Data Collection, Processing, and Normalization

We searched GEO for Illumina 450K and 850K Methylation Array data for undifferentiated human iPSCs and ESCs and selected all studies with at least ten samples, yielding eight studies with a total of 281 female samples, and 190 male samples^38,40–46^. Most of the studies had preprocessed methylation data available. We processed the rest using minfi’s IlluminaNormalization method to mimic the pre-processed data^80^, and combined the 850K and 450K methylation array data into probes that were in common in both arrays. We also removed all of the cross reactive probes as identified by Chen et al in order to exclude ambiguous methylation signals^81^. We further mapped all probes to the hg38 genome assembly using BLAT and removed any that mapped to multiple regions in the genome^82^. After this processing, there were 413,491 probes across the genome, including 9,191 probes on the X.

Mapping the probes to the hg38 genome build also provided updated gene annotations. A gene was assigned to a probe if the probe was within 5kb of the gene. Then the probe was annotated as being either “OutsideTranscript5” (−5000 to −1500 of TSS), “TSS1500” (−1500 to −200 of TSS), “TSS200” (−200 to 0 of TSS), “5_UTR” (in the 5’ UTR of the gene), “Body” (in the body of the gene including exons and introns), “3_UTR” (in the 3’UTR of the gene), and “OutsideTranscript3” (5kb past the end of the gene). All positions were annotated according to a canonical transcript for the gene which was determined to be either the highest transcript support level (TSL) or the highest APPRIS level as annotated by Ensembl^83^. TSL marks the amount of literature support behind a transcript, while APPRIS marks the functional importance of the transcript.

Using the remaining probes, we performed sex prediction using minfi, and removed any lines where the predicted sex did not match the annotated sex. In cases where no annotated sex was available in the originating methylation study, it was obtained from Cellosaurus and compared to the predicted sex^84^. We then performed PCA analyses on the X chromosome probes separately for male and female samples to determine that the data were sufficiently normalized to enable quantitative comparisons between studies (Fig. S1A). Among the female samples, H9 hESCs were overrepresented in the data and tended to separate from most other female hPSCs. To not skew our analysis to differences between H9 and other cell lines, we removed all 60 H9 samples from our analysis (Fig. S1B,C). For the male samples, there 12 of outliers in the PCA that were removed (Fig. S1D,E). This left us with 221 female samples, and 178 male samples.

### Clustering Samples by Variant Probes

In order to calculate which sites had a greater variance in female samples than male samples, we performed a Brown Forsythe (BF) test for each methylation array probe to calculate statistical significance for the difference in variance for the probe between female and male samples. The BF test assigned statistical significance to the differences in variation for each probe between female and male samples. The BF alpha was much more significant for X-probes than for autosomal probes (Fig. S1G). The significantly more variant X-probes also have lower DNAme than the non-sig sites (Fig. S1H). We selected X-linked sites that had a greater variance in the female samples than the male samples (BF p-value ≤ .01), and performed a weighted k-means clustering analysis using the scikit-learn python library on the female samples^85^. The weights for the k-means clustering were based on how many of the cell lines were included in the analysis so that small differences in number of samples per cell line would not skew the k-means calculations. We used the elbow method to determine the appropriate number of clusters to be around six. These clusters were then labeled A-F based on their average X chromosome methylation (most to least methylated). Representative images of how the clusters separated the samples in the PCA is shown in Fig. S1K for a k=4-8.

### Clustering Probes by Progression of XCE

With the samples clustered and ordered, we parsed how the X chromosome probes were changing during XCE. Using the minfi package’s dmpFinder function we found differentially methylated probes between each adjacent sample cluster. Then we divided each CpG site on the X chromosome into one of 11 groups (Fig. 1F, y-axis) based on the most statistically significant transition (smallest p-value ≤ .05), and whether the transition was an increase in methylation or decrease in methylation for the site. These clusters are labeled in figure 1F with a number for the transition they change in, and an arrow to show an increase (↑) or decrease (↓) in methylation. The last cluster (labeled “~”) contains probes that do not pass the p-value threshold (0.05) for all the DMP tests, and thus are not considered to be changing during XCE.

### ATAC-seq Analysis

We aligned the ATAC-seq fastq files using esATAC which serves as a wrapper for AdapterRemoval^86^ for adapter trimming and Bowtie2^87^ for read alignment. MACS2^88^ was used for peak calling, and DiffBind^89^ was used for differential peak analysis. Differential analysis was done using cluster boundaries as for DMP differential methylation analysis.

### Cumulative Demethylation and Calculating Correlation with Xi-specific histone marks

To calculate cumulative methylation changes (grey areas in Fig. 3A), we window-averaged all probes that passed the p-value threshold (≤ .05) in at least one transition and calculated the difference in methylation compared to cluster A moving through the transitions. In contrast, probes that did not pass the threshold had more fluctuant DNAme changes that do not accumulative in successive transitions (Fig. S2B).

We calculated the correlation of the XCE DNAme changes with Xi-specific H3K27me3 and H3K9me3 peaks from^37^. Since these peaks were already window-averaged in the GEO bed files, we decided to window our DNAme data to the same windows (100 kb windows every 50 kb). We used the transition specific changes (colored lines in Fig. 3A) to calculate the Pearson correlation for each transition (Fig. 3C). Similarly, we windowed the differential ATAC peaks (Fig. 3B), and calculated the Pearson correlation for each transition (Fig. 3F).

### Annotating Genes as Escapees

We annotated X chromosome genes as escaping XCI using the DNAme-based escapee calling method presented by Cotton et al^57^ (Fig. S2E-G). We used all male samples, and the female samples from the cluster A to perform this analysis. First, we removed all X chromosome probes with a mean methylation of >= .25 in male samples. To calculate mean DNAme for each gene, we used the TSS for the canonical transcript, and averaged all remaining probes that were situated between −500 and +1500 of the TSS. We calculated a mean TSS methylation separately for the male and cluster A female samples. Assessing the genes defined as the escapee training set, as per Cotton et al, and using the same verification method, we found that only 4/12 their “training set escapees” truly escaped XCI in our dataset (*CA5B, SYAP1, ZFX*, and *ZRSR2*). They had a mean cluster A female methylation of 0.106, and a mean sex difference of 0.02. A gene is classified as an escapee if its female DNAme is less than the mean female DNAme + (3* S.D. female DNAme), and if its sex difference is less than the mean sex difference + (3*S.D. sex difference). Altogether, this approach identified 15 escapee genes (shown in Fig 3A): 12 on the short arm, and 3 on the long arm of the X. We verified the differences in methylation in the promoter and gene body for escapee and subject genes, and found that escapee genes tend to have strongly demethylated promoters and methylation gene bodies on both chromosomes, whereas subject genes have a bimodal distribution of DNAme in both the gene body and the promoter (Fig. S1H).

### Calculating distances to escapee genes

We calculated the distance of eroding DNAme probes from the escapee genes to test whether erosion was emanating from the escapees. For each DMP that was assigned to a transition, we calculated the distance to the nearest escapee gene. We assessed the short and long arms of the X chromosome separately due to the disparity in the number of escapees on each arm. Additionally, we performed two permutation tests to provide randomized distributions against which to compare transition-specific distributions of actual minimal distances. One test randomized the escapee labels across all genes represented by probes, and the other randomized the transition labels for each DMP. Each test was repeated 1000 for each transition and each arm of the chromosome to produce the final distributions (Fig. 3D,E). We repeated this analysis with the differential ATAC peaks instead of the DMPs (Fig 3G,H).

### Expression Analysis

Of the 221 female cell lines we analyzed (after H9 removal), 47 had RNA seq data available. The RNA seq data was downloaded using SRA Toolkit. The FASTQ files were aligned to a hg38 genome assembly (with the PAR region masked on the Y chromosome) using STAR, and reads were counted using HTSeq-count^90,91^.

Differential expression (DE) analysis was performed using DESeq2^92^. We excluded any genes with fewer than 10 counts. DE was performed based on the methylation derived sample clusters and was done individually for each transition rather than considering the cluster assignments as a continuous variable. The differential expression was calculated between each boundary with all the samples. For example, in the transition 1 analysis, we assigned cluster A samples as one group, and clusters B-F as another, then identified differentially expressed genes using DESeq2. For transition 2, we compared clusters A-B to clusters C-F. For transition 3 we compared A-C to D-F. As there were no available expression data for samples in cluster E, we combined transition 4/5 into one and compared clusters A-D to cluster F. By considering all samples, we filtered differentially expressed genes that are not changed in a permanent/consistent manner throughout XCE. We also used DESeq2 to extract VST normalized expression for *XIST, FIRRE, DUSP9, ELK1, REX1*, and *KLF4*, and then ran limma batch correction to adjust for differences in RNA-seq data from different groups. The VST transformation applies a pseudo-log2 transformation on the counts. These VST and batch-corrected values are used for all the expression plots.

### Concordance and Enrichment Analysis

We labeled a gene with concordant methylation and expression as one that had at least one methylation array probe changing in the same transition as the expression and having the opposite direction of change as the expression (i.e. decreasing methylation for genes increasing in expression, and vice versa). We further allowed a gene to be considered concordant if it had a differential ATAC peak with +/− 3kb of the gene changing in the same transition as the change in expression (all concordant genes are shown in Fig. S3C). To check for concordance between expression and ATAC-seq, we simply checked for differential ATAC peaks to be within 3kb of the gene. The genes that were either concordant with the DNAme or with the ATAC peaks are shown in figure S3C.

Using these transition-specific concordant gene sets, we calculated ontology term enrichments in each transition for X chromosome genes, and autosomal genes. Using the clusterProfiler R package, we queried the following gene sets: Disease Ontology, DisGeNET, GO, KEGG, MeSH gendoo, MSigDB, Network of Cancer Gene, Reactome, and WikiPathways. For the cancer gene counts (Fig. 4F), we used the COSMIC Cancer Gene Census to count how many cancer genes of the various types were in each transition^79^.

### Telescope HERV Analysis

We used Telescope^67^ to enumerate the HERV expression for each STAR aligned bam files from the expression analysis. Then we followed the same DESeq2 pipeline from the expression analysis to calculate differential HERV expression between clusters and to obtain the log2-transformed VST+batch-corrected expression for each sample.

### ATAC HERV Overlap Analysis

Using bedtools fisher, we compared the overlap for each repeat from RepeatMasker and our differential ATAC-peaks from each transition. We used the right-sided p-value to select overlaps that were enriched in our differential ATAC-peaks. Figure 4J plots the significant overlaps for HERVs that were also differentially expressed in Transition 4/5 (Fig. 4I).

## SUPPLEMENTARY FIGURE LEGENDS

**Figure S1: DNAme data processing to distinguish clusters along XCE**

(A) Schematic of filtering and processing pipeline for DNA methylation data.

(B) and (C) PCAs of female samples on X-probes before and after H9 removal, respectively. Plots are colored by different properties of the samples as indicated by the legends.

(D) and (E) PCAs of male samples on X-probes before and after outlier removal, respectively. The plots are colored by different properties of the samples as indicated by the legends.

(F) Global mean methylation for female and male samples (N.S. = not significant).

(G) Brown-Forsythe test p-values for autosomal probes and X-probes showing that X-probes are significantly more variant between the male and female samples than autosomal probes.

(H) Distribution of ß-values for X-probes that are more variant in female samples (Sig Sites), and those that are not (Non-sig-sites).

(I) and (J) Relationship between the mean X chromosome methylation and mean methylation of non-sig-x-probes or sig-x-probes, respectively. The line shows the linear regression, and “corr” is the Pearson correlation.

(K) Plots of the elbow method to find the appropriate number of clusters for the k-means clustering of female samples.

**Figure S2: DNAme dynamics during XCE distinguish escapees and reactivated genes**

(A) The percent composition of probes in each probe grouping from Figure 2A, as described in the methods. Probe totals for each group are listed to the right.

(B) Cumulative changes through each transition for X-linked probes with high sex-variance but without consistent direction across transitions (“ ~ “ symbol), plotted as in Fig. 3A.

(C) and (D) VST+batch-corrected expression for *REX1* and *KLF4*, respectively. *REX1* has 2 outliers hidden in the dot plot to ease boxplot visualization.

(E-G) Identification of escapees using the Cotton et al method. “Sex Delta” panels show the difference in methylation for a gene between male and female samples. (E) methylation values of the initial Cotton, et al. trainee genes, (F) as G after removal of initial trainee genes that fail cutoffs (as in Cotton et al), and (G) final escapee gene set after true designation is determined and the escapees are identified.

(H) Distribution of β-values of gene body and promoter probes for escapees and genes subject to XCI.

(I) Cumulative probability of CpG demethylation in each transition as a function of distance to escapee genes on the whole X chromosome.

(J) as in K, exclusively for CpGs on the short arm.

(K) as in K, exclusively for CpGs on the long arm.

(L) Cumulative probability of ATAC-peak openings in each transition as a function of distance to escapee genes on the whole X chromosome.

(M) as in L, exclusively for ATAC-peaks on the short arm.

(N) as in L, exclusively for ATAC-peaks on the long arm.

**Figure S3: Global changes in DNAme and gene expression**

(A) Chromosome-resolved DMPs assigned to specific transitions by p-value. Total counts for autosomal and X chromosome probes are provided, with direction of change indicated (blue = loss of DNAme, and red = gain of DNAme). The transparency is on a continuous scale based on the β-differential.

(B) Chromosome-resolved differential ATAC peaks assigned to specific transitions by FDR. Total counts are provided as for (A) (blue = closing ATAC peak, red = opening ATAC peak). The transparency is on a continuous scale based on the log2(fold change) of the ATAC peaks.

(C) Chromosome-resolved differentially expressed genes concordant with either DMP (A) or differential ATAC peaks (B). Total counts are provided as for (A) (blue = decreasing expression, and red = increasing expression). The transparency is on a continuous scale based on the log2(fold change) of the expression.

**Figure S4: Autosomal DNAme changes relative to *DUSP9* reactivation impact naïve markers**

(A) Relationship of *DUSP9* promoter methylation to autosomal methylation in female samples. Clusters E and F are emphasized to highlight the change in global and *DUSP9* methylation in the last transition.

(B) Change in methylation for genes that are differentially expressed in Transition 4/5 and are differentially expressed in naïve stem cells^64^.

(C) DNAme heatmap of *ELK1* probes across clusters A-F.

(D) *ELK1* (VST+batch-corrected/log2-transformed) expression across clusters A-F.

## Notes

### Competing Interest Statement

The authors have declared no competing interest.

