## Supplementary figures and images for "Contiguous Erosion of the Inactive X in Human Pluripotency Concludes With Global DNA Hypomethylation"

### Supplemental Figures

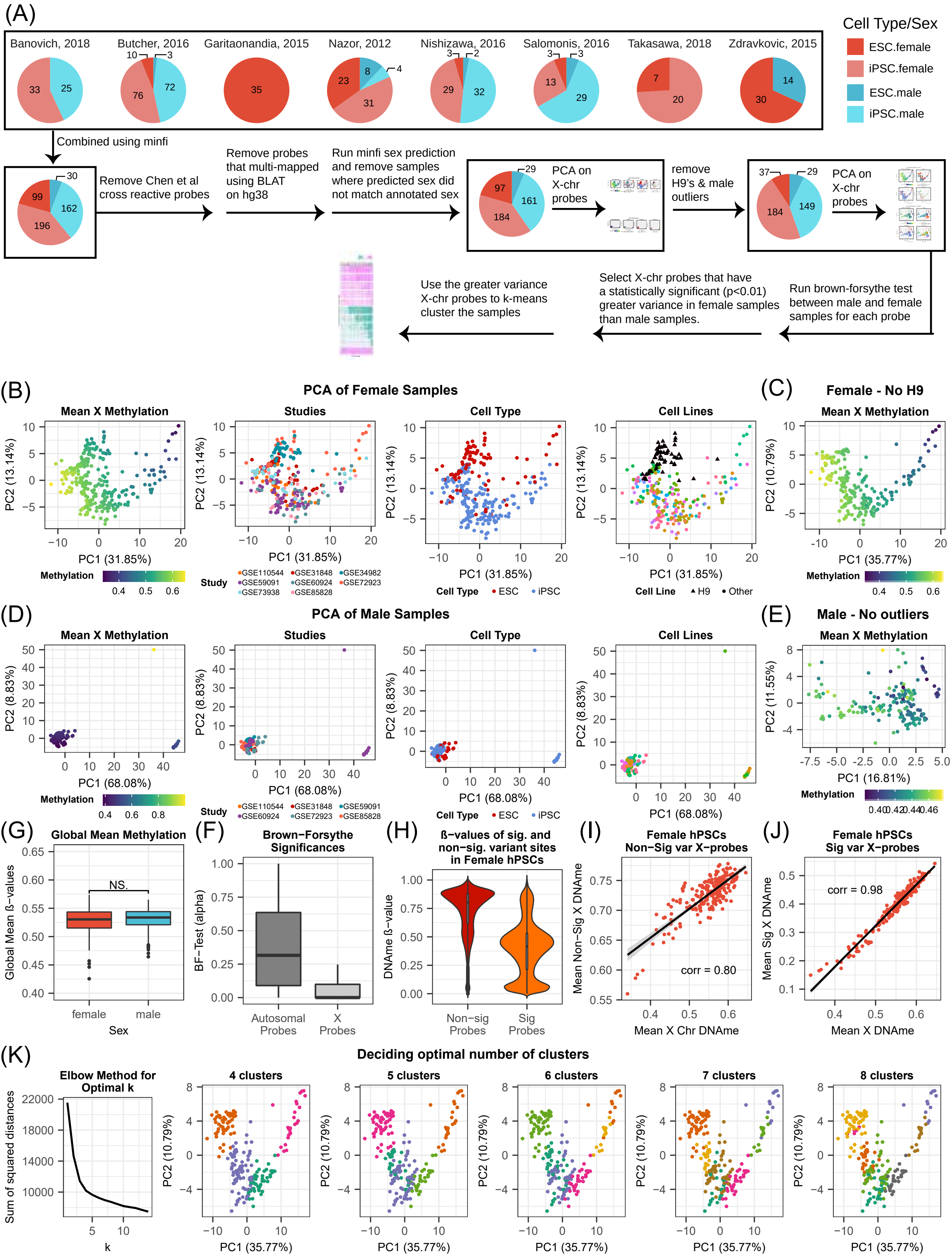

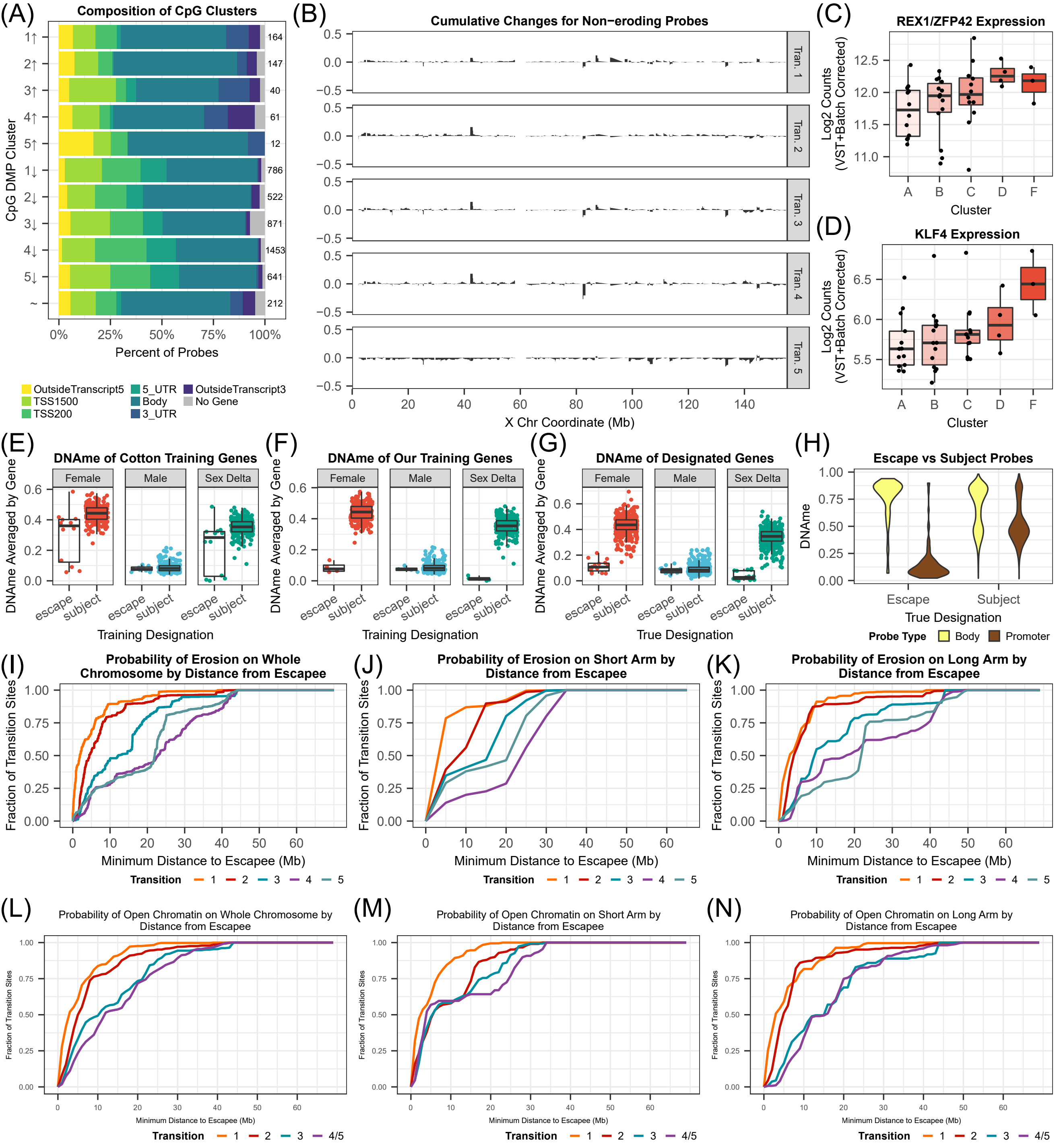

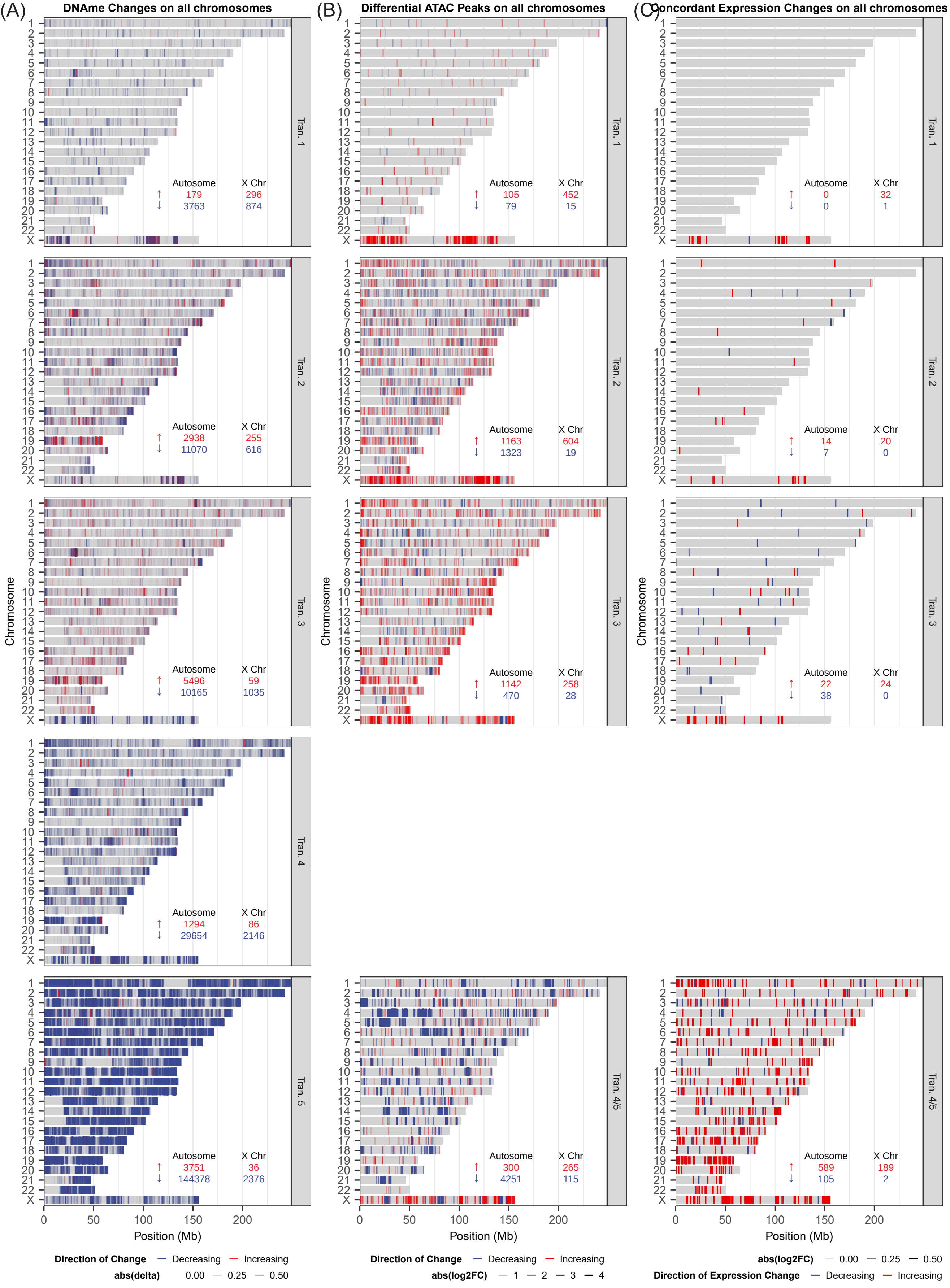

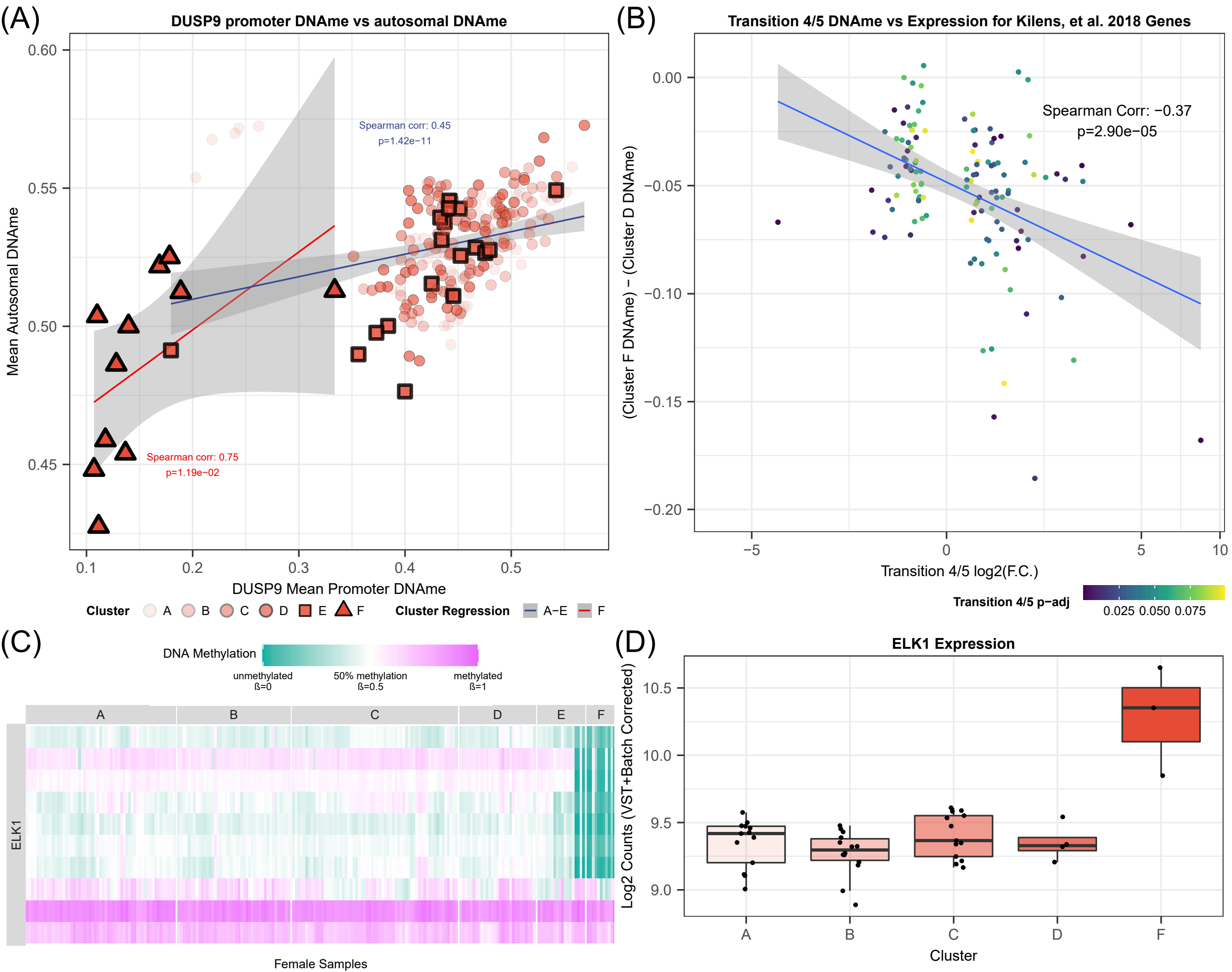
